# Host Adaptation Predisposes *Pseudomonas aeruginosa* to Type VI Secretion System-Mediated Predation by the *Burkholderia cepacia* Complex

**DOI:** 10.1101/2020.04.10.036012

**Authors:** Andrew I Perault, Courtney E Chandler, David A Rasko, Robert K Ernst, Matthew C Wolfgang, Peggy A Cotter

## Abstract

*Pseudomonas aeruginosa* (*Pa*) and *Burkholderia cepacia* complex (Bcc) species are opportunistic lung pathogens of individuals with cystic fibrosis (CF). While *Pa* can initiate long-term infections in younger CF patients, Bcc infections only arise in teenagers and adults. Both *Pa* and Bcc use type VI secretion systems (T6SS) to mediate interbacterial competition. Here, we show that *Pa* isolates from teenage/adult CF patients, but not those from young CF patients, are outcompeted by the epidemic Bcc isolate *Burkholderia cenocepacia* strain AU1054 (*Bc*AU1054) in a T6SS-dependent manner. The genomes of susceptible *Pa* isolates harbor T6SS-abrogating mutations, the repair of which, in some cases, rendered the isolates resistant. Moreover, seven of eight Bcc strains outcompeted *Pa* strains isolated from the same patients. Our findings suggest that certain mutations that arise as *Pa* adapts to the CF lung abrogate T6SS activity, making *Pa* and its human host susceptible to potentially fatal Bcc superinfection.

## INTRODUCTION

The respiratory tracts of individuals suffering from the genetic disorder cystic fibrosis (CF) are hospitable environments for microorganisms, and thus CF patients harbor complex, dynamic microbial communities in their airways that can include opportunistic pathogens (Carmody *et al*., 2015; Filkins and O’Toole, 2015; Lipuma, 2010; J. Zhao *et al*., 2012). *Pseudomonas aeruginosa* and certain members of the *Burkholderia cepacia* complex (Bcc), a taxonomic group containing at least 17 *Burkholderia* spp. (Salsgiver *et al*., 2016), cause devastating infections in CF patients (Govan and Deretic, 1996; Lipuma, 2010; Mahenthiralingam *et al*., 2005). While *P. aeruginosa* infects young CF patients and is the most common opportunistic CF pathogen by early adulthood, Bcc infections are less common and, for unknown reasons, limited to older CF patients, typically teenagers and adults (Cystic Fibrosis Foundation, 2019). Unlike other CF pathogens, Bcc strains are more frequently associated with person-to-person spread (Biddick *et al*., 2003; Chen *et al*., 2001; Govan *et al*., 1993) and can progress to a fatal necrotizing pneumonia and bacteremia termed “cepacia syndrome” (Isles *et al*., 1984; Lipuma, 2010). While *P. aeruginosa*-Bcc co-infections occur within CF patients, the two pathogens do not colocalize. *P. aeruginosa* is mostly found extracellular in the airway lumen and Bcc within phagocytes; however, the *P. aeruginosa* burden in co-infected patients tends to be lower than in patients infected by *P. aeruginosa* alone (Schwab *et al*., 2014).

Given the polymicrobial nature of the CF respiratory tract, interbacterial interactions likely occur in these tissues and may influence disease progression (Bisht *et al*., 2020; Filkins and O’Toole, 2015; O’Brien and Fothergill, 2017; Peters *et al*., 2012). Interbacterial competition is hypothesized to be one of the strongest determinants of ecology and evolution within polymicrobial communities (Foster and Bell, 2012). A prevalent and well-understood mechanism of interbacterial competition is the type VI secretion system (T6SS) (Alteri and Mobley, 2016; Russell *et al*., 2014), which is predicted to be present in ∼25% of Gram-negative bacteria (Boyer *et al*., 2009). T6SSs use a bacteriophage-like mechanism to deliver effector proteins directly into target bacterial or eukaryotic cells (Basler *et al*., 2012; Hachani *et al*., 2016; Hood *et al*., 2010; Pukatzki *et al*., 2007). Antibacterial T6SS effectors disrupt diverse biological processes within target cells, and cognate immunity proteins protect T6SS-producing cells from autotoxicity (Ahmad *et al*., 2019; Russell *et al*., 2014; Ting *et al*., 2018). Type VI secretion (T6S) has been studied in the Bcc pathogen *Burkholderia cenocepacia* strain J2315 (*Bc*J2315), which produces a T6SS that is important for infection of macrophages and influences the immune response to this pathogen (Aubert *et al*., 2015; 2016; Hunt *et al*., 2004; Rosales-Reyes *et al*., 2012). Recent bioinformatic analysis has detected T6SS-encoding genes throughout the Bcc, with one system (referred to as T6SS-1) predominating; however, several species encode multiple T6SSs (Spiewak *et al*., 2019). The *B. cenocepacia* strain H111 T6SS was shown to have modest antibacterial activity (Spiewak *et al*., 2019), though the potential antibacterial role of T6SSs in other Bcc pathogens remains unknown.

*P. aeruginosa* produces three separate T6SSs (the H1-, H2-, and H3-T6SSs), and while both the H1- and H2-T6SSs are antibacterial weapons, the H1-T6SS is the stronger mediator of interbacterial competition (Allsopp *et al*., 2017; Hood *et al*., 2010; Russell *et al*., 2011). The *P. aeruginosa* H1-T6SS is under intricate regulation at both the post-transcriptional and post-translational level. Phosphorelay through the GacSA two-component system activates T6SS protein production via the regulatory small RNAs (sRNAs) RsmY and RsmZ, which relieve RsmA-mediated repression of T6SS transcript translation (Goodman *et al*., 2004; 2009; Lapouge *et al*., 2008; Moscoso *et al*., 2011; Ventre *et al*., 2006). Moreover, a threonine phosphorylation pathway regulates T6SS assembly and function via signal transduction through the membrane-associated TagQRST proteins, Fha1, and the kinase and phosphatase PpkA and PppA, respectively (Basler *et al*., 2013; Casabona *et al*., 2013; Hsu *et al*., 2009; Mougous *et al*., 2007).

*P. aeruginosa* undergoes dramatic evolution within the CF respiratory tract to transition to a chronic infection lifestyle (Folkesson *et al*., 2012; Winstanley *et al*., 2016), and evolutionary analyses have detected mutations in *gacS*/*gacA* as well as T6SS structural genes (Bartell *et al*., 2019; Kordes *et al*., 2019; Marvig *et al*., 2015). Given Bcc infections establish later in the lives of CF patients compared to *P. aeruginosa* infections, we hypothesized that host adaptation by *P. aeruginosa* may open the door to subsequent Bcc infections if the resident *P. aeruginosa* community loses T6SS activity. In this study, we conducted experiments to test this hypothesis.

## RESULTS

### The *Bc*AU1054 T6SS mediates interbacterial competition

We selected *Burkholderia cenocepacia* strain AU1054 (*Bc*AU1054), which was isolated from the bloodstream of a CF patient who succumbed to the infection, for our studies (Chen *et al*., 2001; Grigoriev *et al*., 2012). The *Bc*AU1054 genome encodes a predicted T6SS on chromosome 1 (BCEN_RS13060, *tssM*, through BCEN_RS13170, *tssL*) (**Figure 1A**). Presumably due to errors during the whole genome sequencing of this strain (assembly GCA_000014085.1), multiple genes in this region were annotated as pseudogenes. We PCR amplified and sequenced these regions and found that each gene is intact, encoding a full-length protein (**Table S1)**. Compared to the T6SS-encoding cluster of *B. cenocepacia* strain J2315 (*Bc*J2315), the *Bc*AU1054 T6SS gene cluster contains an additional region (BCEN_RS13075 (*tai1*) through BCEN_RS13110 (*vgrG1*)) that includes two predicted effector-immunity (E-I)-encoding gene pairs (*tae1*-*tai1* and *tle1*-*tli1*) (**Figure S1**). Immediately 3’ to the *Bc*AU1054 core cluster (with the 5’ to 3’ direction corresponding to the sequence numbering in **Figure 1A**) is another additional region containing *vgrG2*, the predicted E-I-encoding pair *tne1*-*tni1*, and several genes predicted to encode proteins involved in phage or other mobile genetic elements.

**Figure 1.**
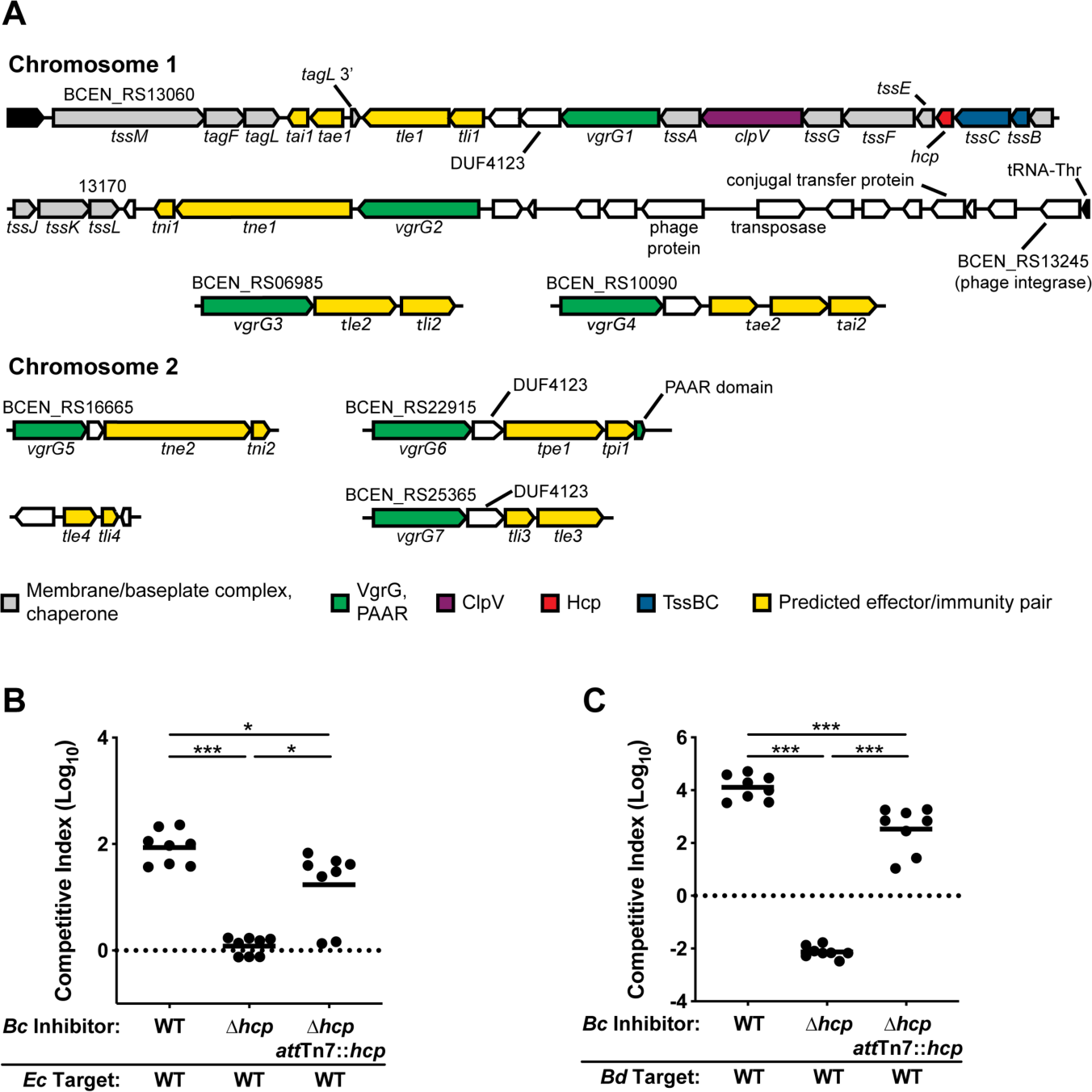
The *Bc*AU1054 T6SS mediates interbacterial competition. (A) Core cluster (top two lines) and accessory genes encoding the *Bc*AU1054 T6SS and bioinformatically-predicted effector-immunity (E-I) pairs (see **Table S2**). The legend indicates function of protein products. White ORFs encode DUF4123 T6SS adapter proteins, or proteins not known to be associated with the T6SS. (B and C) Interbacterial competition experiments between inhibitor *Bc*AU1054 and target *E. coli* (B) and *Bd*AU0158 (C). WT, Δ*hcp*, and Δ*hcp att*Tn7::*hcp Bc*AU1054 inhibitor strains used in each. Circles represent individual co-cultures from two biological replicates, each with four technical replicates. Solid horizontal lines represent mean log_10_ C.I. values. Dotted horizontal lines (log_10_ C.I. = 0) indicate no competitive advantage for either inhibitor or target. \**P*<0.05, \*\**P*<0.0005, Mann-Whitney test.

To identify additional genes with the potential to encode E-I pairs, we first analyzed genes located near the seven annotated *vgrG* genes (**Figure 1A**). VgrG proteins, along with PAAR-repeat proteins, form the puncturing tip of the T6SS needle and typically associate with effectors encoded by nearby genes (Pukatzki *et al*., 2007; Russell *et al*., 2014; Shneider *et al*., 2013). Based on previous nomenclature (Russell *et al*., 2014), we named predicted cell membrane-degrading effectors Tle for <u>T</u>6SS lipase <u>e</u>ffector, nucleic acid-degrading effectors Tne for <u>T</u>6SS <u>n</u>uclease <u>e</u>ffector, cell wall-degrading effectors Tae for <u>T</u>6SS <u>a</u>midase <u>e</u>ffector, and used Tpe for the <u>T</u>6SS <u>p</u>ore-forming <u>e</u>ffector. We named cognate immunity proteins Tli, Tni, Tai, and Tpi. Three of the predicted E-I-encoding gene pairs (*tle1*-*tli1, tle3*-*tli3*, and *tpe1*-*tpi1*) are found near ORFs encoding proteins with the domain of unknown function (DUF) 4123, which is a highly conserved chaperone domain for T6SS effectors (Liang *et al*., 2015) (**Figure 1A**). Protein secondary structure analysis using Phyre2 (Kelley *et al*., 2015) and HHpred (Zimmermann *et al*., 2018) predicted antibacterial enzymatic activities for all potential effectors (**Table S2**). Tae1 and Tle4 are not encoded by genes closely associated with *vgrG* genes, but have predicted antibacterial activities. A duplication of the 3’ end of *tagL* flanking *tae1*-*tai1* (**Figure 1A**) suggests these predicted E-I-encoding genes inserted into the *Bc*AU1054 T6SS core cluster via a transposon, and secondary structure analysis predicts a glycosyl hydrolase domain in Tae1. We identified the *tle4*-*tli4* gene pair by searching for DUFs shared among E-I-encoding genes, as DUF3304 is only present in the *Bc*AU1054 genome within *tli1, tli3*, and *tli4* (**Table S2**).

To determine whether the *Bc*AU1054 T6SS mediates interbacterial competition, we generated an unmarked, in-frame deletion mutation in *hcp*, which encodes the inner tube protein of the T6SS apparatus. Over 5 h co-culture, *Bc*AU1054 outcompeted *Escherichia coli* DH5α by ∼2-logs, whereas *Bc*AU1054 Δ*hcp* had no competitive advantage (**Figure 1B**). *Bc*AU1054 also had a T6SS-dependent competitive advantage over another Bcc pathogen, *Burkholderia dolosa* strain AU0158 (*Bd*AU0158), as the parental strain outcompeted *Bd*AU0158 by ∼4-logs over 5 h, whereas *Bc*AU1054 Δ*hcp* was outcompeted by *Bd*AU0158 (**Figure 1C**). Ectopic expression of *hcp* from the *att*Tn*7* site of the genome partially restored the ability of *Bc*AU1054 Δ*hcp* to outcompete *E. coli* DH5α and *Bd*AU0158 (**Figures 1B** and **1C**). Importantly, *Bc*AU1054 Δ*hcp* did not have a growth defect compared to the parental strain, suggesting growth rate differences were not a factor determining competitive fitness (**Figure S2**). The *Bc*AU1054 T6SS, therefore, is a potent antibacterial weapon capable of killing competitor bacteria.

### At least five *Bc*AU1054 T6SS effectors mediate interbacterial competition

To determine which predicted effectors are involved in T6SS-mediated interbacterial competition by *Bc*AU1054, we generated nine mutants, each containing an unmarked, in-frame deletion mutation in one of the predicted E-I-encoding gene pairs. We screened these mutants by engineering them to produce green fluorescent protein (GFP) and co-culturing them, individually, with either wild-type (WT) or Δ*hcp Bc*AU1054 strains for ∼20 h, and then measuring GFP fluorescence intensity and the OD_600_ of the co-cultures. For four of the mutants (Δ*tle1*Δ*tli1*, Δ*tne1*Δ*tni1*, Δ*tne2*Δ*tni2*, and Δ*tpe1*Δ*tpi1*), the GFP/OD_600_ values for co-cultures with WT *Bc*AU1054 were about half of what they were for co-cultures with *Bc*AU1054 Δ*hcp*, indicating these mutants were outcompeted in a T6SS-dependent manner, presumably because they lack the immunity protein that is specific for the cognate toxic effector (**Figure S3**). To measure competition quantitatively, we then competed each of these four mutants against WT and Δ*hcp Bc*AU1054 strains. In each case, the E-I deletion mutant was outcompeted by its parental strain in a T6SS-dependent manner (**Figure 2A**). Ectopic expression of the cognate immunity gene in each mutant rescued it from T6SS-mediated killing by the parental *Bc*AU1054 strain (**Figure 2A**), providing evidence that Tle1-Tli1, Tne1-Tni1, Tne2-Tni2, and Tpe1-Tpi1 are true antibacterial E-I pairs associated with the *Bc*AU1054 T6SS.

**Figure 2.**
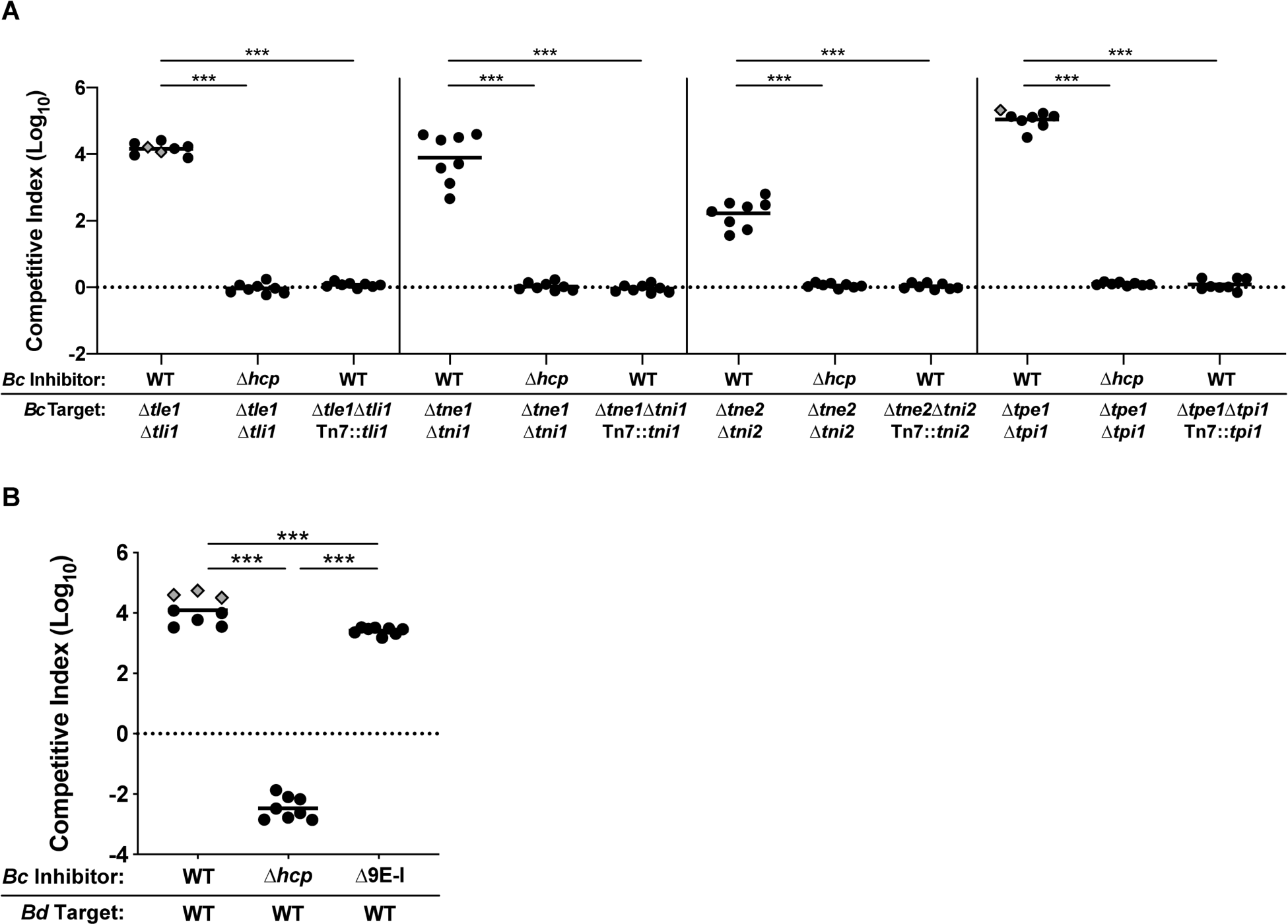
At least five *Bc*AU1054 T6SS effectors mediate interbacterial competition. (A) Interbacterial competition experiments between WT and Δ*hcp Bc*AU1054 inhibitor strains and *Bc*AU1054 mutants lacking E-I-encoding genes, including mutants complemented with cognate immunity genes. Probable E-I pairs identified by screen shown in **Figure S3**. (B) Interbacterial competition experiments between WT, Δ*hcp*, and Δ9E-I *Bc*AU1054 inhibitor strains and *Bd*AU0158 target. For (A and B), circles/diamonds represent individual co-cultures from two biological replicates, each with four technical replicates. Grey-filled diamonds represent competitions from which no target cells were recovered. Solid horizontal lines represent mean log_10_ C.I. values. Dotted horizontal lines (log_10_ C.I. = 0) indicate no competitive advantage for either inhibitor or target. \*\*\**P*<0.0005, Mann-Whitney test.

To determine if there are additional E-I-encoding gene pairs in *Bc*AU1054, we generated a mutant lacking all nine predicted E-I-encoding gene pairs (*Bc*AU1054 Δ9E-I) and assessed its ability to outcompete target bacteria. This nonuple mutant outcompeted *Bd*AU0158 by ∼3.5-logs (slightly less than WT *Bc*AU1054) (**Figure 2B**), indicating that at least one more effector delivered by the *Bc*AU1054 T6SS exists.

### Susceptibility of *P. aeruginosa* CF isolates to the *Bc*AU1054 T6SS correlates with patient age

Since Bcc pathogens are known to establish infections in the polymicrobial CF respiratory tract, we next sought to determine whether the *Bc*AU1054 T6SS targets the prevalent CF pathogen *P. aeruginosa*. In competition experiments against *P. aeruginosa* reference strain PAO1, PAO1 outcompeted both WT and Δ*hcp* strains of *Bc*AU1054, though showed a slightly (∼0.5-log) greater ability to outcompete T6SS-active than T6SS-inactive *Bc*AU1054 (**Figure 3A**). These results are consistent with two theories on the regulation of H1-T6SS activity by PAO1: T6SS-dueling (Basler *et al*., 2013), in which the PAO1 H1-T6SS only deploys following antagonism by a neighboring cell, and the *P. aeruginosa* response to antagonism (PARA) (LeRoux *et al*., 2015), in which PAO1 activates aggressive behaviors, like T6SS activity, following detection of kin cell lysates. The ability of PAO1 to outcompete *Bc*AU1054 was solely dependent on the H1-T6SS, as a transposon insertion in *vipA1*, which encodes a necessary structural protein of the H1-T6SS (Basler *et al*., 2013), rendered PAO1 susceptible to being outcompeted by *Bc*AU1054 (**Figure 3A**).

**Figure 3.**
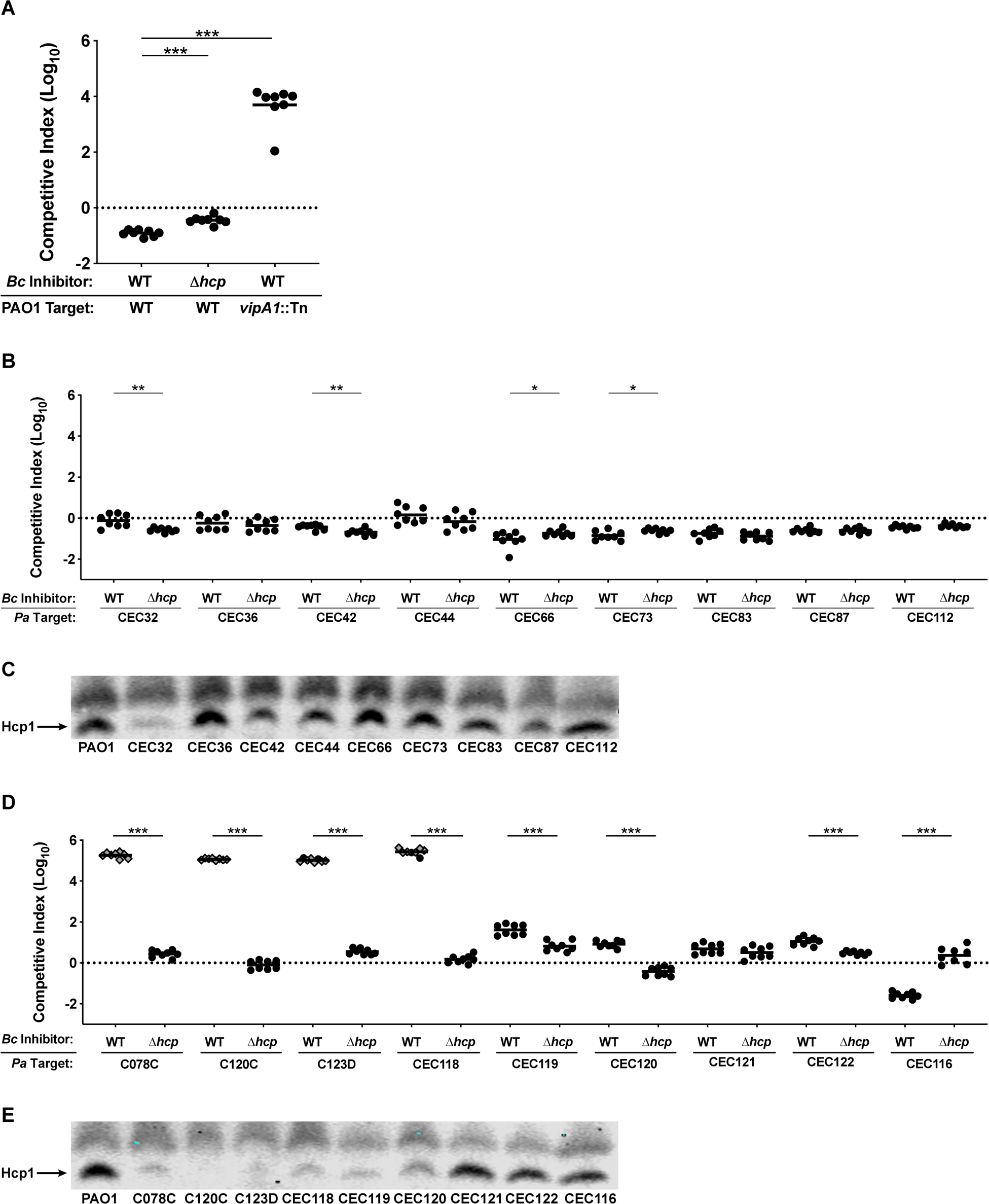
Susceptibility of *P. aeruginosa* CF isolates to the *Bc*AU1054 T6SS correlates with patient age. (A) Interbacterial competition experiments between WT and Δ*hcp Bc*AU1054 inhibitor strains and WT and *vipA1*::Tn PAO1 target strains. (B) Interbacterial competition experiments between WT and Δ*hcp Bc*AU1054 inhibitor strains and *P. aeruginosa* infant/child CF isolate targets. (C) Immunoblotting for Hcp1 production by PAO1 and *P. aeruginosa* infant/child CF isolates. (D) Interbacterial competition experiments between WT and Δ*hcp Bc*AU1054 inhibitor strains and *P. aeruginosa* teenage/adult CF isolate targets. (E) Immunoblotting for Hcp1 production by PAO1 and *P. aeruginosa* teenage/adult CF isolates. For (A, B, and D), circles/diamonds represent individual co-cultures from two biological replicates, each with four technical replicates. Grey-filled diamonds represent competitions from which no target cells were recovered. Solid horizontal lines represent mean log_10_ C.I. values. Dotted horizontal lines (log_10_ C.I. = 0) indicate no competitive advantage for either inhibitor or target. \**P*<0.05, \*\**P*<0.005, \*\*\**P*<0.0005, Mann-Whitney test. For (C and E), non-specific band above Hcp1 serves as loading control. Blots are representative of at least two experiments.

PAO1 was originally isolated from a wound infection and has undergone decades of laboratory passage and diversification (Chandler *et al*., 2019; Holloway, 1955; Holloway and Morgan, 1986; Klockgether *et al*., 2010). We therefore reasoned that PAO1 may not accurately represent potential T6SS-mediated interactions between Bcc pathogens and *P. aeruginosa* strains relevant to CF infection. To address this concern, we used collections of *P. aeruginosa* strains isolated from CF patients (Burns *et al*., 2001; Rosenfeld *et al*., 2001). *Bc*AU1054 did not outcompete any of the *P. aeruginosa* strains isolated from infants or young children (three years old or younger) and was often slightly outcompeted by these *P. aeruginosa* strains (**Figure 3B**). By contrast, *Bc*AU1054 had the striking ability to outcompete nearly half of the *P. aeruginosa* strains isolated from teenagers and adults (11-31 years old) in a T6SS-dependent manner, oftentimes efficiently enough to prevent recovery of any *P. aeruginosa* from the co-cultures (**Figure 3D**). We also determined if the nonuple E-I deletion mutant of *Bc*AU1054 could outcompete the susceptible *P. aeruginosa* strains. Although *Bc*AU1054 Δ9E-I was strongly outcompeted by PAO1, it retained a competitive advantage against C078C, C120C, C123D, and CEC118 (**Figure S4**). Intriguingly, C120C was less susceptible to killing by *Bc*AU1054 Δ9E-I than were C078C, C123D, and CEC118, (**Figure S4**) suggesting C120C is less sensitive to the unidentified effector(s) associated with the *Bc*AU1054 T6SS, and that *Bc*AU1054 T6SS effectors exhibit target strain-specific variability in toxicity. The ability of *Bc*AU1054 to efficiently kill *P. aeruginosa* from older CF patients correlates with the clinical presentation of Bcc infections, which do not occur in young CF patients and solely arise in teenagers and adults (Cystic Fibrosis Foundation, 2019).

### Host-adapted *P. aeruginosa* isolates that are sensitive to the *Bc*AU1054 T6SS harbor T6SS-abrogating mutations

We sequenced the genomes of all *P. aeruginosa* clinical isolates (not from co-infections) used in our study. Of the four T6SS-susceptible teenage/adult isolates (C078C, C120C, C123D, and CEC118), three contain mutations in the *gacS* or *gacA* genes, which encode a two-component system required for T6SS protein production (**Table 1**) (Goodman *et al*., 2004; Marden *et al*., 2013; Moscoso *et al*., 2011). C078C contains a missense mutation in *gacS* (*gacS*_G1715A_), resulting in the variant protein GacS_G572D_. C123D contains a large genomic deletion spanning the *gacS* gene and the nearby *pirRSA* genes, which encode a siderophore iron-acquisition system (Ghysels *et al*., 2005). C120C contains a premature stop codon in *gacA* (*gacA*_C349T_). The C078C genome also contains a premature stop codon in *pppA* (*pppA*_G111A_). PppA is a post-translational regulator of H1-T6SS activity (Mougous *et al*., 2007), and is required for efficient T6SS-mediated competition (Basler *et al*., 2013). CEC118 has a small deletion in *fha1* (*fha1*_Δ404-424_) resulting in the loss of seven amino acid residues from Fha1, another post-translational regulator of H1-T6SS activity (Mougous *et al*., 2007).

**Table 1.**
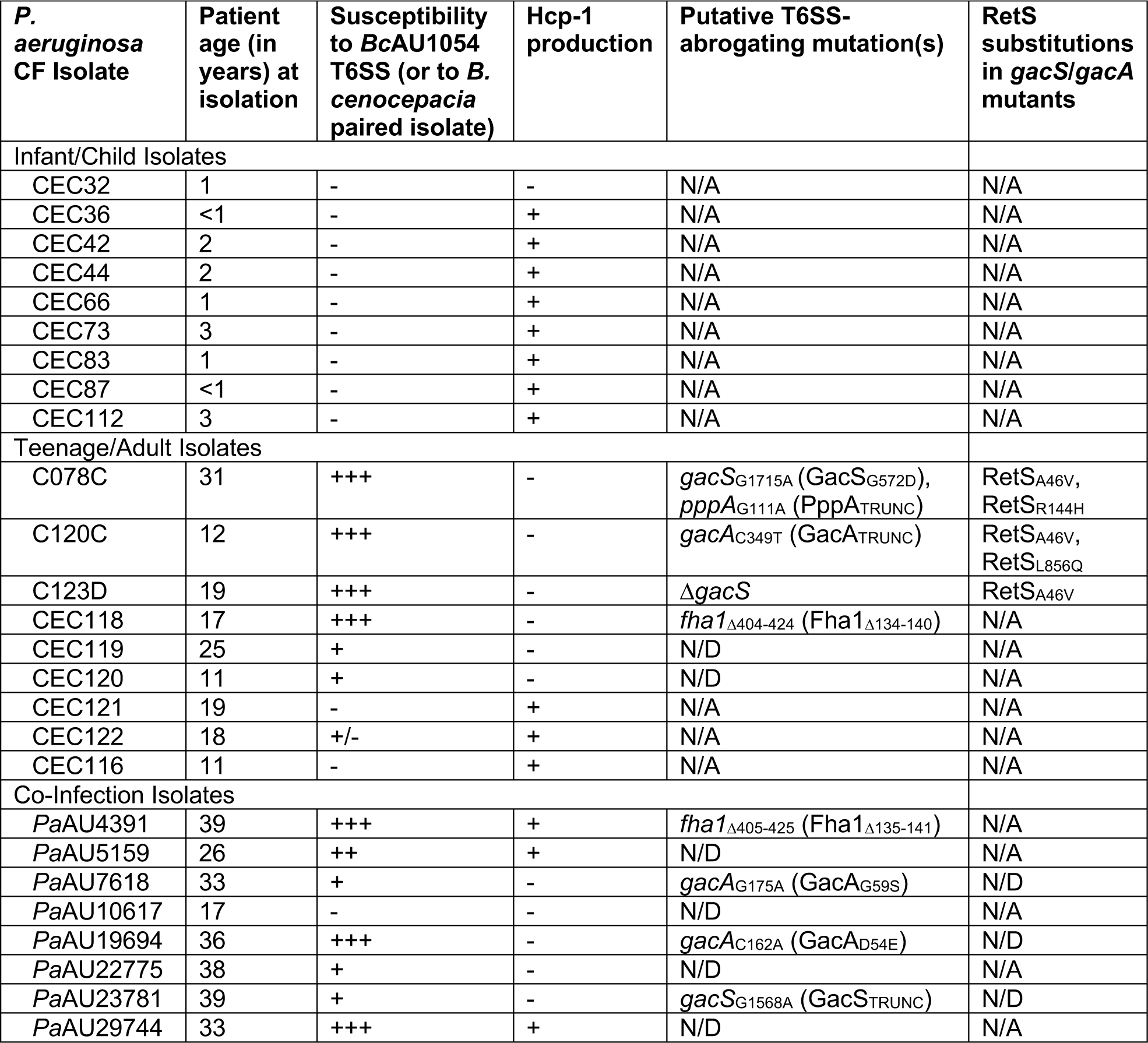
Competition sensitivity, Hcp1 production, and putative T6SS-abrogating mutations of *P. aeruginosa* CF isolates used in this study. Isolates grouped into infant/child isolates, teenage/adult isolates, and co-infection isolates, with patient age at isolation provided. For susceptibility to competition against *B. cenocepacia* and Hcp1 production status, see **Figures 3** and **6**. N/A, not applicable. N/D, not determined.

To investigate if the ability of *Bc*AU1054 to outcompete *P. aeruginosa* strains isolated from teenage/adult CF patients correlates with a loss of H1-T6SS activity in the *P. aeruginosa* strains, we assessed production of Hcp1, the major subunit protein of the H1-T6SS inner tube, during growth on agar. Every *P. aeruginosa* isolate that was outcompeted by the *Bc*AU1054 T6SS showed either negligible or diminished Hcp1 production compared to PAO1 (**Figures 3D** and **3E**). The isolates CEC121, CEC122, and CEC116, which were not outcompeted by the *Bc*AU1054 T6SS, produced Hcp1 at levels similar to PAO1 (**Figure 3E**). Conversely, every *P. aeruginosa* strain isolated from an infant or young child except CEC32 produced Hcp1 at or near levels similar to PAO1 (**Figure 3C**), which correlates with their resistance to T6SS-dependent outcompetition by *Bc*AU1054 (**Figure 3B**).

### Restoration of H1-T6SS protein production can rescue host-adapted *P. aeruginosa* from T6SS-mediated elimination by *Bc*AU1054

Phosphorelay through the GacSA two-component system activates production of the sRNAs RsmY and RsmZ, which are required for T6SS protein production by *P. aeruginosa* (Goodman *et al*., 2009; 2004; Lapouge *et al*., 2008; Moscoso *et al*., 2011; Ventre *et al*., 2006). To determine if lack of *gacS*/*gacA* function is responsible for susceptibility of the *P. aeruginosa* strains with mutations in these genes, we introduced a plasmid (pJN-*rsmZ*) into C120C, C123D, and C078C to express *rsmZ* (induced by arabinose) independent of the GacSA phosphorelay (Intile *et al*., 2014; Janssen *et al*., 2018). We also introduced the vector backbone (pJN105) into these strains as a negative control. In competitions against *Bc*AU1054 on agar containing 0.1% arabinose, C120C pJN-*rsmZ* was rescued from T6SS-mediated elimination by *Bc*AU1054, while C120C harboring the pJN105 was not (**Figure 4A**). Likewise, C120C pJN-*rsmZ*, but not C120C pJN105, produced Hcp1 when grown under inducing conditions (**Figure 4D**). C123D pJN-*rsmZ* (**Figure 4B**) and C078C pJN-*rsmZ* (**Figure 4C**) were still strongly outcompeted by the *Bc*AU1054 T6SS, and pJN-*rsmZ* did not promote Hcp1 production by these isolates (**Figure 4D**). Because C078C also contains a premature stop codon in *pppA*, we delivered the WT *pppA* gene under control of a constitutive promoter to the *att*Tn*7* site, and also introduced pJN-*rsmZ* into this strain, but expression of these genes failed to rescue this strain from T6SS-mediated elimination by *Bc*AU1054 (**Figures 4E** and **4F**). Lastly, we delivered the WT *fha1* gene to the *att*Tn7 site of CEC118, as this isolate has a truncated *fha1* (*fha1*_Δ404-424_), but constitutive expression of full-length *fha1* did not rescue CEC118 from being outcompeted by *Bc*AU1054 (**Figure 4G**). It is not surprising that constitutive expression of full-length *pppA* and full-length *fha1* did not rescue C078C and CEC118, respectively, as both isolates were defective for Hcp1 production (**Figure 3E**); moreover, the natively-produced truncated PppA and Fha1 variants could act as dominant negatives in these strains.

**Figure 4.**
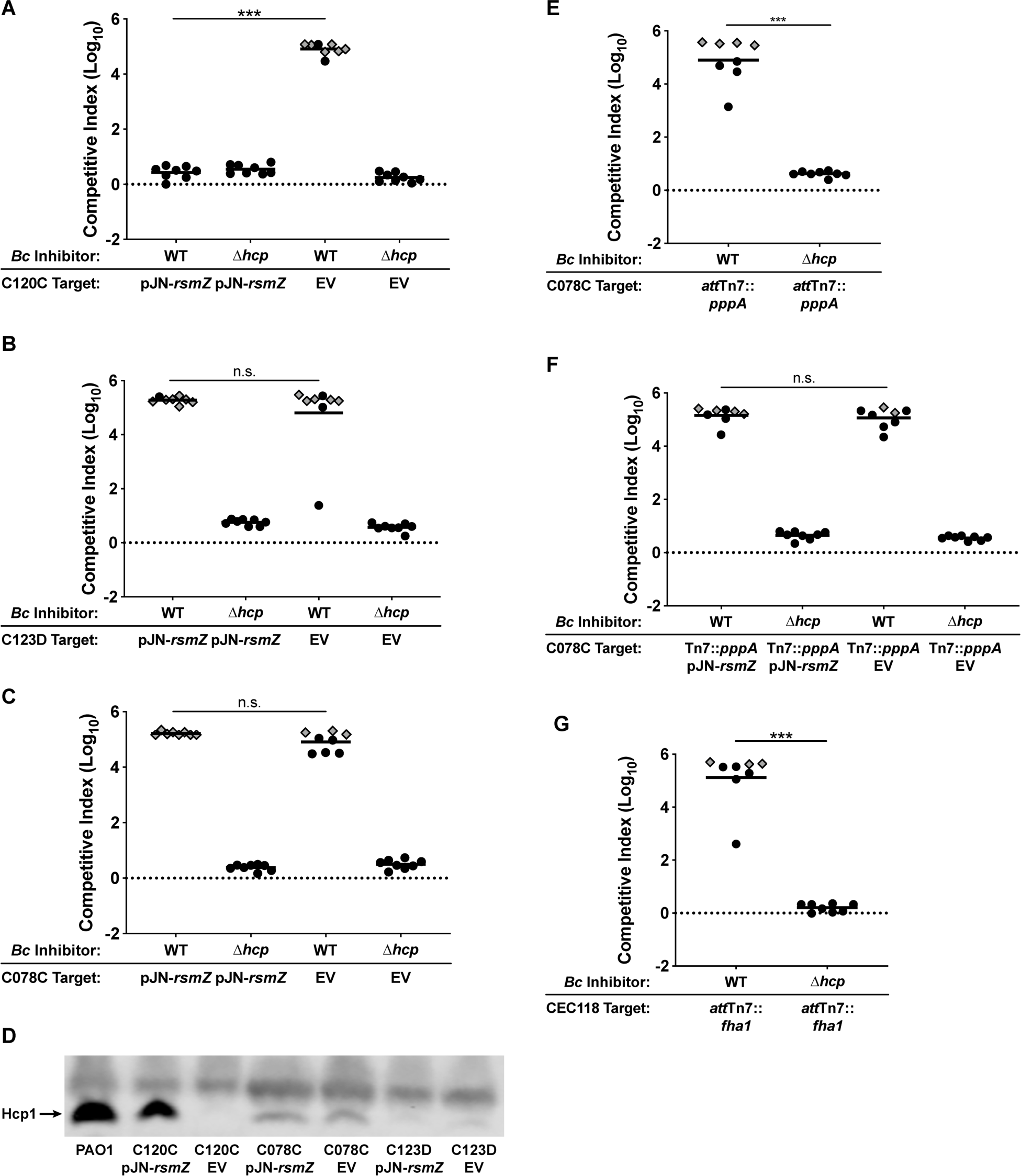
Restoration of H1-T6SS protein production can rescue host-adapted *P. aeruginosa* from T6SS-mediated elimination by *Bc*AU1054. (A, B, and C) Interbacterial competition experiments between WT and Δ*hcp Bc*AU1054 inhibitor strains and *P. aeruginosa* teenage/adult CF isolates C120C (A), C123D (B), and C078C (C) harboring the pJN-*rsmZ* and pJN105 (EV) plasmids. Competitions conducted on agar containing 0.1% arabinose. (D) Immunoblotting for Hcp1 production by C120C, C078C, and C123D isolates harboring pJN-*rsmZ* and pJN105 (EV) during growth on agar containing 0.1% arabinose, as well as Hcp1 production by PAO1 for comparison. Non-specific band above Hcp1 serves as loading control. The blot is representative of at least two experiments. (E) Interbacterial competition experiments between WT and Δ*hcp Bc*AU1054 inhibitor strains and C078C *att*Tn7::*pppA* target strain. (F) Interbacterial competition experiments between WT and Δ*hcp Bc*AU1054 inhibitor strains and C078C *att*Tn7::*pppA* target strains harboring pJN-*rsmZ* and pJN105 (EV). Competitions conducted on agar containing 0.1% arabinose. (G) Interbacterial competition experiments between WT and Δ*hcp Bc*AU1054 inhibitor strains and CEC118 *att*Tn7::*fha1* target strain. For (A, B, C, E, F, and G), circles/diamonds represent individual co-cultures from two biological replicates, each with four technical replicates. Grey-filled diamonds represent competitions from which no target cells were recovered. Solid horizontal lines represent mean log_10_ C.I. values. Dotted horizontal lines (log_10_ C.I. = 0) indicate no competitive advantage for either inhibitor or target. n.s.=not significant, \*\*\**P*<0.0005, Mann-Whitney test.

### Additional Bcc pathogens kill host-adapted *P. aeruginosa* in a T6SS-dependent manner

To investigate whether T6SS-mediated killing of host-adapted *P. aeruginosa* is a common feature of Bcc strains, we used *Burkholderia multivorans* strain CGD2M (*Bm*CGD2M) and *Bd*AU0158, which encode predicted T6SS-1 systems. *Bm*CGD2M encodes one additional predicted T6SS, and *Bd*AU0158 encodes two additional predicted T6SSs. We generated plasmid disruption mutations in the *tssC1* genes of these strains’ T6SS-1 clusters (*Bm*CGD2M *tssC1*::pAP82 and *Bd*AU0158 *tssC1*::pAP83), and competed these mutants and the parental strains against *P. aeruginosa* strains PAO1 and C078C. *Bm*CGD2M had a slight (∼1-log) T6SS-dependent competitive advantage against PAO1, and strongly outcompeted the *P. aeruginosa* host-adapted isolate C078C using its T6SS (**Figure 5A**). Although *Bd*AU0158 was outcompeted by PAO1, possibly via T6SS-dueling or PARA on the part of PAO1, *Bd*AU0158 outcompeted C078C by ∼3-logs in a T6SS-dependent manner (**Figure 5B**). These data suggest T6S may provide many Bcc pathogens a competitive advantage against host-adapted *P. aeruginosa*.

**Figure 5.**
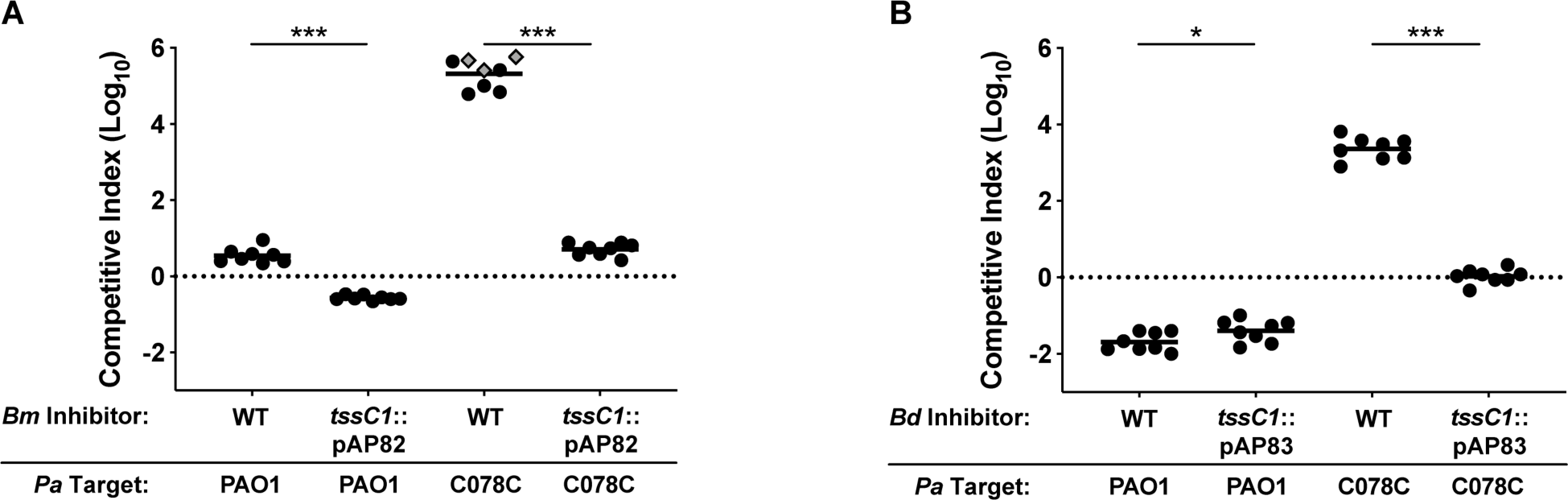
Additional Bcc pathogens kill host-adapted *P. aeruginosa* in a T6SS-dependent manner. (A) Interbacterial competition experiments between WT and *tssC1*::pAP82 *Bm*CGD2M inhibitor strains and PAO1 and C078C target strains. (B) Interbacterial competition experiments between WT and *tssC1*::pAP83 *Bd*AU0158 inhibitor strains and PAO1 and C078C target strains. For (A and B), circles/diamonds represent individual co-cultures from two biological replicates, each with four technical replicates. Grey-filled diamonds represent competitions from which no target cells were recovered. Solid horizontal lines represent mean log_10_ C.I. values. Dotted horizontal lines (log_10_ C.I. = 0) indicate no competitive advantage for either inhibitor or target. \**P*<0.05, \*\*\**P*<0.0005, Mann-Whitney test.

### *B. cenocepacia* isolates from CF patients with concurrent *P. aeruginosa* infections outcompete their paired *P. aeruginosa* isolates under T6SS-permissive conditions

Our data, together with data from other groups (Bartell *et al*., 2019; Kordes *et al*., 2019; Marvig *et al*., 2015), suggest T6SS-abrogating mutations can accumulate as *P. aeruginosa* evolves within the CF respiratory tract to transition to a chronic infection lifestyle, and that patients infected by T6SS-null *P. aeruginosa* may be susceptible to Bcc superinfections. To explore this hypothesis further, we acquired eight *B. cenocepacia*-*P. aeruginosa* coinfection pairs, each isolated from a separate CF patient with concurrent *B. cenocepacia* and *P. aeruginosa* infections. During co-culture on agar, seven of the eight *B. cenocepacia* isolates outcompeted their paired *P. aeruginosa* isolate by as little as ∼1-log (*Bc*AU7523 vs. *Pa*AU7618) or as great as ∼4-logs (*Bc*AU29704 vs. *Pa*AU29744) (**Figure 6A**). Since genetic manipulation of recent human isolates is often not possible, we took advantage of the fact that growth in shaking liquid cultures is non-permissive for T6SS-mediated competition (Hood *et al*., 2010; Majerczyk *et al*., 2016; Russell *et al*., 2011; Speare *et al*., 2020), likely because cells are not in contact long enough to allow T6SS effector delivery to target cells. The competitive advantages of *B. cenocepacia* isolates over their *P. aeruginosa* paired isolates dropped dramatically during shaking liquid growth compared to growth on agar (**Figure 6B**); while some isolates (*Bc*AU7523, *Bc*AU22760) only experienced an ∼1-log decrease in competitive index, others (*Bc*AU4392, *Bc*AU19695, *Bc*AU29704) experienced an ∼3-log decrease in competitive index. In fact, liquid growth provided two *P. aeruginosa* isolates (*Pa*AU7618 and *Pa*AU19694) a competitive advantage over their paired *B. cenocepacia* isolates (**Figure 6B**).

**Figure 6.**
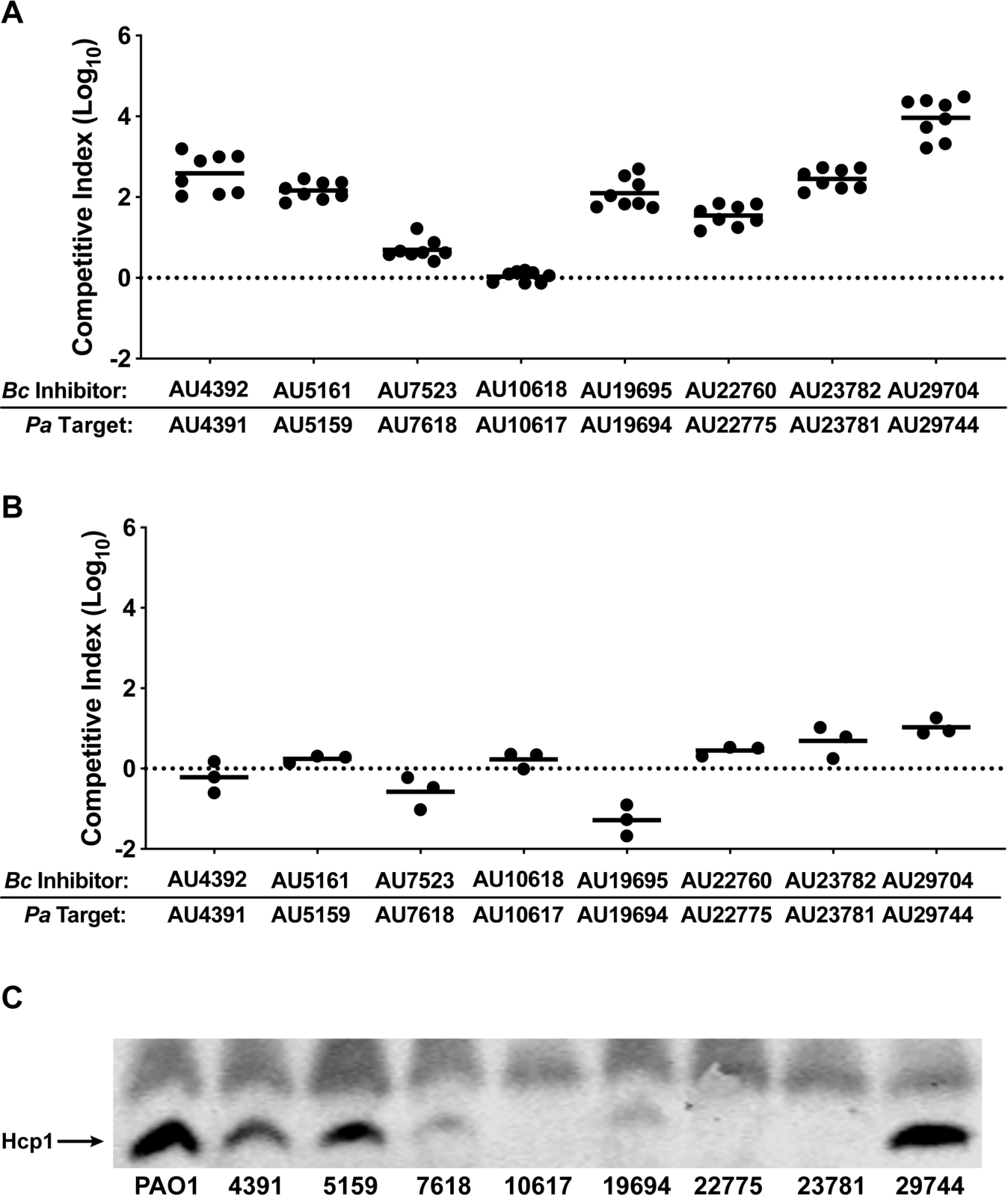
*B. cenocepacia* isolates from CF patients with concurrent *P. aeruginosa* infections outcompete their paired *P. aeruginosa* isolates under T6SS-permissive conditions. (A and B) Interbacterial competition experiments between *B. cenocepacia*-*P. aeruginosa* co-infection isolate pairs on agar (A) and in shaking liquid culture (B). For agar competitions in (A), circles represent individual co-cultures from two biological replicates, each with four technical replicates. For shaking liquid competitions in (B), circles represent individual co-cultures from three biological replicates. Solid horizontal lines represent mean log_10_ C.I. values. Dotted horizontal lines (log_10_ C.I. = 0) indicate no competitive advantage for either inhibitor or target. (C) Immunoblotting for Hcp1 production by *P. aeruginosa* co-infection isolates, as well as Hcp1 production by PAO1 for comparison. Non-specific band above Hcp1 serves as loading control. The blot is representative of at least two experiments.

To identify potential genetic explanations for the competitive disadvantages of the *P. aeruginosa* co-infection isolates, we PCR-amplified the genes involved in H1-T6SS production/activity that are mutated in the *P. aeruginosa* strains isolated from adults for which we have whole-genome sequence information (**Table 1**) and sequenced these PCR products. Three isolates contain mutations in *gacS* or *gacA* (*Pa*AU7618 contains a *gacA*_G175A_ mutation resulting in GacA_G59S_, *Pa*AU19694 contains a *gacA*_C162A_ mutation resulting in GacA_D54E_, and *Pa*AU23781 contains a premature stop codon in *gacS* (*gacS*_G1568A_)) (**Table 1**). *Pa*AU4391 contains a small deletion in *fha1* (*fha1*_Δ405-425_) that is nearly identical to the mutation in CEC118 (*fha1*_Δ404-424_) (**Table 1**). Western blotting showed negligible or diminished Hcp1 production by five *P. aeruginosa* co-infection isolates (*Pa*AU7618, *Pa*AU10617, *Pa*AU19694, *Pa*AU22775, and *Pa*AU23781) compared to PAO1 (**Figure 6C**). *Pa*AU4391, *Pa*AU5159, and *Pa*AU29744 produced Hcp1 at levels similar to PAO1 (**Figure 6C**), though *Pa*AU4391 has an *fha1*_Δ405-425_ mutation that may abrogate T6SS activity without affecting protein production. *Pa*AU5159 and *Pa*AU29744 may harbor mutations in other genes important for post-translational regulation of T6SS activity. These results suggest that Bcc pathogens may only be able to invade a *P. aeruginosa*-colonized CF respiratory tract if the *P. aeruginosa* population, or at least a subpopulation, has evolved to lose T6SS activity.

## DISCUSSION

Although the propensity of Bcc pathogens to infect only older CF patients, and to cause superinfections in those colonized with *P. aeruginosa*, has been appreciated for many years (Folescu *et al*., 2015; McCloskey *et al*., 2001; Whiteford *et al*., 1995), the underlying reasons for this apparent selectivity are unknown. During our investigation of T6S in the Bcc, we found that none of the *P. aeruginosa* strains in our study that were isolated from infant or child CF patients were susceptible to T6SS-mediated killing by *Bc*AU1054. Conversely, almost half of the strains isolated from teenage and adult CF patients were outcompeted by *Bc*AU1054 in a T6SS-dependent manner, with susceptible *P. aeruginosa* strains often being completely eliminated from the co-cultures. Additional Bcc pathogens (*Bm*CGD2M and *Bd*AU0158) also efficiently outcompeted susceptible *P. aeruginosa* strains via T6SS activity, and seven of eight *B. cenocepacia* strains from patients with concurrent *P. aeruginosa* infections outcompeted their paired *P. aeruginosa* strains under conditions promoting T6SS-mediated interactions. These data suggest that one reason Bcc pathogens are restricted to infecting older CF patients is because only in these patients are resident *P. aeruginosa* susceptible to T6SS-mediated competition by Bcc bacteria.

We found that differential susceptibility of *P. aeruginosa* strains to T6SS-mediated competition by Bcc pathogens depends on T6SS functionality in *P. aeruginosa*. Disruption of *vipA1* to inactivate the H1-T6SS in PAO1 converted it from being resistant to T6SS-mediated competition by *Bc*AU1054 to being outcompeted by four logs. For the susceptible *P. aeruginosa* strains isolated from teenagers or adults, we found that all harbor mutations predicted to abrogate production and/or function of their T6SSs, all failed to produce substantial amounts of Hcp1, and in one strain, elimination by *Bc*AU1054 was prevented by activating production of its T6SS proteins. Consistent with these observations, *B. cenocepacia* isolates strongly outcompeted their co-isolated *P. aeruginosa* strains only when the bacteria were co-cultured on a solid surface (conducive to contact-dependent interactions) and not when co-cultured in liquid medium. Furthermore, the *P. aeruginosa* co-infection isolates were typically deficient in Hcp1 production. These data indicate that, at least for the *P. aeruginosa* strains studied here, the main factor in determining susceptibility to T6SS-mediated competition by Bcc bacteria is whether *P. aeruginosa* produces a functional T6SS.

The T6S-abrogating mutations we identified in *P. aeruginosa* CF isolates in this study fell into two classes: those in genes encoding post-translational regulators of T6SS activity (*pppA* and *fha1*), and those in genes encoding the GacSA two-component regulatory system. Fha1 is required for the initial assembly of the T6S apparatus, whereas PppA is required for disassembly of apparatuses and recycling of T6SS proteins into new apparatuses. Mutations in *pppA* or *fha1* are expected to prevent efficient T6SS activity without affecting production of individual T6SS components (Basler *et al*., 2013; Mougous *et al*., 2007). Consistent with this expectation, Hcp1 was detectable in *Pa*AU4391, which contains a small deletion in *fha1*, but this co-infection isolate was outcompeted by its paired *B. cenocepacia* isolate. By contrast, mutations in *gacA* or *gacS* are expected to prevent production of the entire T6S apparatus. GacS is one of four hybrid sensor kinases that controls phosphorylation, and hence activation, of the GacA response regulator. LadS functions with GacS to activate GacA, while RetS blocks GacS activity, thereby inhibiting GacA activation (Chambonnier *et al*., 2016; Goodman *et al*., 2009). The PA1611-encoded sensor kinase promotes GacA activation by relieving RetS inhibition of GacS (Kong *et al*., 2013). When active, GacA induces production of two sRNAs, RsmY and RsmZ, which bind to, and prevent activity of, RsmA, a pleiotropic global regulator that impedes translation of many target genes (Brencic and Lory, 2009; Brencic *et al*., 2009). When not inhibited by RsmY or RsmZ, RsmA activity results in production of factors associated with acute infection (e.g., flagella, type III secretion, type IV pili) and lack of production of factors and phenotypes associated with chronic infection (e.g., exopolysaccharide production, biofilm, T6S). The RetS/PA1611/LadS/GacSA signaling pathway is therefore considered to function as a switch between acute and chronic infection modes (Balasubramanian *et al*., 2013; Goodman *et al*., 2009; 2004).

While there is evidence that the genes encoding the RetS/PA1611/LadS/GacSA signaling pathway are intact when *P. aeruginosa* establishes infection initially in the CF lung, mutations arise in *retS* within some strains over time (e.g., 11/36 clone types in the 2015 Marvig *et al*. study), and, at least for those studied, all *retS*-mutated strains acquire subsequent mutations in *gacS*/*gacA* or *rsmA* (Bartell *et al*., 2019; Marvig *et al*., 2015). Our data are consistent with these reports, as three out of nine teenage/adult *P. aeruginosa* isolates used in our study were *gacS*/*gacA* mutants and also contained nonsynonymous *retS* mutations (these isolates were also defective in Hcp1 production). Three out of eight of the *P. aeruginosa* co-infection isolates contained *gacS*/*gacA* mutations and did not produce Hcp1; their *retS* statuses are unknown. Thus, there appears to be a selection for lack of GacSA activity following mutation of *retS* within the CF respiratory tract, and we envisage that this selection could be either T6S-independent or T6S-dependent; a Gac-regulated target other than T6S may drive this selection, with loss of T6S being simply a consequence of Gac inactivation, or T6SS activity itself could be what is selected against. We and others (Kordes *et al*., 2019; Marvig *et al*., 2015) have detected mutations in genes encoding proteins specific for T6SS assembly and function in *P. aeruginosa* strains isolated from older CF patients, supporting the hypothesis that T6S may be disadvantageous to *P. aeruginosa* during chronic infection in the CF lung.

Why might it be beneficial for *P. aeruginosa* to lose T6SS activity in later stages of host colonization? Given the polymicrobial nature of the CF respiratory tract, it is reasonable to hypothesize that maintaining a potent antibacterial weapon like the T6SS would be beneficial. However, *P. aeruginosa* T6SS proteins are immunogenic (Mougous *et al*., 2006), and avoiding the host immune response could be equally, or more, beneficial. Additionally, production of T6SSs is energetically costly, and while T6SS-mediated competition may be worth the cost during early stages of infection, it may be beneficial to stop producing these structures once *P. aeruginosa* has established its niche. As indicated by the proportion of reads in metagenomic samples, *P. aeruginosa* can constitute over 90% of all bacterial cells within the airways of certain CF patients (Carmody *et al*., 2015; 2013; J. Zhao *et al*., 2012). Under these conditions, T6S-mediated inter-species competition should not be required. Loss of T6S by predominant bacterial strains colonizing humans has been shown with gut resident *Bacteroides* spp., as T6SS-proficient *Bacteroides* are more prevalent in the unstable infant gut microbiota than they are in adult gut microbiota where individual *Bacteroides* spp. or strains predominate (Verster *et al*., 2017). One might expect that similar selective pressures would also act on Bcc during CF infection. However, the *B. cenocepacia* T6SS is required for murine infection (Hunt *et al*., 2004), and at least some Bcc strains produce a T6S-dependent effector (TecA) that promotes intracellular survival within macrophages (Aubert *et al*., 2016; 2015; Rosales-Reyes *et al*., 2012), suggesting there is a strong selective advantage for Bcc pathogens to remain T6SS-active during infection of the CF respiratory tract.

Although *P. aeruginosa* and Bcc bacteria ultimately colonize different sites in the CF airways (Schwab *et al*., 2014), Bcc pathogens must traverse the lumen, where *P. aeruginosa* can exist in large populations, before invading host cells. Therefore, transient Bcc-*P. aeruginosa* interactions likely occur, and our data support the hypothesis that the outcome of these interactions depends on the T6S proficiency of the resident *P. aeruginosa. P. aeruginosa* populations within individual CF patients exhibit genotypic and phenotypic diversity across different regions of the respiratory tract (Jorth *et al*., 2015), and thus Bcc bacteria may only need to interact with a subpopulation of *P. aeruginosa* that has lost T6SS activity in order to initiate an infection and invade host cells. Experiments using animal models and human microbiome analyses have shown that T6SS-mediated competition occurs within mammalian intestines (M. C. Anderson *et al*., 2017; Sana *et al*., 2016; Verster *et al*., 2017; Wexler *et al*., 2016; W. Zhao *et al*., 2018), though it is not known whether such interactions occur in the CF respiratory tract. These questions would be better addressed with animal models of CF disease. Unfortunately, a dearth of robust, efficient animal models for chronic bacterial infections has inhibited progress in the understanding of these infections (Fisher *et al*., 2011; Kukavica-Ibrulj and Levesque, 2008; Semaniakou *et al*., 2018).

While our data are consistent with T6SS-mediated competition between Bcc pathogens and *P. aeruginosa* playing a role in susceptibility of older CF patients to the Bcc, we hypothesize that additional factors play important roles in preventing Bcc infections in infants and young children. *Staphylococcus aureus* is the most prevalent pathogen of young CF patients (Cystic Fibrosis Foundation, 2019), and *S. aureus* colonization could preclude infection by Bcc pathogens. Additionally, changes in the immune response, physiology, and/or nutritional environment of the CF respiratory tract over time could cause these tissues to be more hospitable to Bcc pathogens later in the lives of CF patients compared to those in infants and children. CF patients are also often on antibiotic regimens to treat opportunistic infections, and regular use of antibiotics may promote Bcc pathogen colonization of older patients. Other unknown factors could also be at play.

In our studies, competition mediated by the T6SS-1 provided strong competitive advantages to three Bcc pathogens (*Bc*AU1054, *Bm*CGD2M, and *Bd*AU0158) against host-adapted *P. aeruginosa*. Gene clusters encoding the T6SS-1 are prevalent throughout the Bcc (Spiewak *et al*., 2019), suggesting that T6SS-mediated killing of host-adapted *P. aeruginosa* may be a common asset of Bcc pathogens. The role of additional T6SSs in Bcc pathogens remains unknown, but it appears that interbacterial antagonism is mostly mediated by the T6SS-1, at least under the conditions used in this study. Our investigation of the *Bc*AU1054 T6SS revealed four *bona fide* antibacterial E-I pairs; however, our bioinformatic prediction of E-I pair-encoding genes missed at least one gene pair, as *Bc*AU1054 Δ9E-I maintained a strong competitive advantage against *Bd*AU0158. The unidentified effector(s) is/are not encoded by gene(s) near *vgrG* genes, nor are there shared domains between the effectors we identified and the unidentified effector(s), suggesting the unidentified effector(s) may be members of an uncharacterized class of T6SS toxins. Future studies will identify the full repertoire of *Bc*AU1054 T6SS E-I pairs. Our screening for antibacterial effectors and follow-up competition experiments were specific to intrastrain antagonism (*Bc*AU1054 vs. *Bc*AU1054) under one condition (LSLB agar at 37°C). The predicted E-I pairs that our screen suggested were not important for intrastrain competition may be important for interstrain/interspecies competition or competition under different conditions (e.g., temperature, salt, pH); similar conditional efficiency has been demonstrated for *P. aeruginosa* T6SS effectors (LaCourse *et al*., 2018). Supporting this hypothesis, *Bc*AU1054 Δ9E-I outcompeted T6SS-null *P. aeruginosa* teenage/adult isolates to varying degrees, suggesting the additional, unidentified effector(s) have prey cell-specific activity.

There is growing appreciation for the genotypic and phenotypic diversity of *P. aeruginosa* within the CF respiratory tract (Folkesson *et al*., 2012; Jorth *et al*., 2015; Winstanley *et al*., 2016). Although reference strains are powerful tools for studying bacterial pathogens, they do not always perfectly represent the strains currently infecting humans. Our investigations illuminate differences between PAO1 and recently collected *P. aeruginosa* CF isolates specific to T6SS-mediated competition against Bcc pathogens, as well as demonstrate varying abilities of *P. aeruginosa* CF isolates to compete against *Bc*AU1054. Our data support a model in which resident *P. aeruginosa* populations must evolve to lose T6SS activity in order for Bcc pathogens to colonize the CF respiratory tract. If true, not only is the Bcc T6SS an important colonization factor, but assessing the T6S potential of resident *P. aeruginosa* could predict susceptibility of CF patients to deadly Bcc superinfections.

## METHOD DETAILS

### Bacterial strains and growth conditions

All bacterial strains in this study were cultured in low salt lysogeny broth (LSLB: 10 g/L tryptone, 5 g/L yeast extract, 5 g/L sodium chloride) or on LSLB agar (1.5% agar). Antibiotics to select for *Burkholderia* strains were used at the following concentrations, when applicable: 30 µg/mL gentamicin, 250 µg/mL kanamycin, 50 µg/mL trimethoprim, 40 µg/mL tetracycline. Antibiotics to select for *P. aeruginosa* strains were used at the following concentrations, when applicable: 20 or 35 µg/mL chloramphenicol, 20 µg/mL nalidixic acid, 50µg/mL trimethoprim, 75 µg/mL gentamicin, 40 µg/mL tetracycline. 20 µg/mL nalidixic acid was used to select for *E. coli* DH5α, when applicable.

### Genetic manipulations

*E. coli* strain RHO3 was used to conjugate plasmids into *Burkholderia* spp. and *P. aeruginosa*. The pEXKm5 allelic exchange vector (López *et al*., 2009) was used to generate unmarked, in-frame deletion mutations in *Bc*AU1054. Briefly, ∼500 nucleotides 5’ to and including the first three codons of the gene to be deleted were fused to ∼500 nucleotides 3’ to and including the last three codons of the gene by overlap extension PCR and cloned into pEXKm5. Following selection of *Bc*AU1054 merodiploids with the plasmids integrated into the chromosome, cells were grown for 4 h in YT broth (10 g/L yeast extract, 10 g/L tryptone) at 37°C with aeration, subcultured 1:1000 in fresh YT broth, and grown overnight at 37°C with aeration. After overnight growth, cells that lost the cointegrated plasmid following the second homologous recombination step were selected on YT agar (1.5% agar) containing 25% sucrose and 100 µg/mL 5-bromo-4-chloro-3-indoxyl-β-D-glucuronide (X-Gluc). Deletion mutants were screened for by PCR and verified by sequencing regions spanning the deletions.

The pUC18T-mini-Tn7T suite of plasmids (Choi *et al*., 2005) was used to deliver antibiotic resistance gene cassettes to the *att*Tn7 sites of *Bc*AU1054 and *P. aeruginosa*. The trimethoprim resistance-conferring plasmid pUC18T-mini-Tn7T-Tp was generated in this study by restriction digesting out *dhfRII* from pUC18T-mini-Tn7-Tp-P_S12_-mCherry (LeRoux *et al*., 2012) using MscI and NcoI and ligating into digested pUC18T-mini-Tn7T-Km (Choi *et al*., 2005) lacking *nptII* (the kanamycin resistance-conferring gene). pUC18-miniTn7-*kan*-*gfp* (Norris *et al*., 2010) was used to generate GFP-producing *Bc*AU1054 E-I deletion mutants. Complemented *Bc*AU1054 mutant strains (with either *hcp* or T6SS immunity-encoding genes) were generated by PCR amplifying the genes of interest and cloning the sequences into pUCS12Km (M. S. Anderson *et al*., 2012). *Bc*AU1054 strains constitutively expressing *lacZ* were generated using pECG10 (M. S. Anderson *et al*., 2012). *P. aeruginosa* isolates C078C and CEC118 were complemented with *pppA* and *fha1*, respectively, by cloning these sequences into pUCS12Km, digesting out the genes and upstream constitutive promoter P_S12_, and cloning these fragments into pUC18T-mini-Tn7T-Tet (M. S. Anderson *et al*., 2012). For all pUC18T-mini-Tn7T-based cassette delivery to the *att*Tn7 sites of *Bc*AU1054 and *P. aeruginosa*, the transposase-encoding pTNS3 helper plasmid was used in triparental conjugation. *Bm*CGD2M *tssC1*::pAP82 and *Bd*AU0158 *tssC1*::pAP83 were generated by cloning ∼500 internal nucleotides of the *tssC1* genes into pUC18T-mini-Tn7T-Km, conjugating the plasmids into *Bm*CGD2M and *Bd*AU1058, and selecting for plasmid cointegrants on kanamycin.

### Interbacterial competition experiments

All competition experiments were conducted for 5 h on LSLB agar at 37°C, with an ∼1:1 starting cell ratio of inhibitor and target strains, unless stated otherwise. Cells were collected from overnight liquid cultures, centrifuged for 2 min at 15,000 rpm, washed in 1X phosphate buffered saline (PBS), diluted to an OD_600_ of 1.0, and equal volumes of inhibitor and target cells were mixed. For *Bc*AU1054 vs. *E. coli* DH5α competitions, *Bc*AU1054 1.0 OD_600_ cell suspensions were diluted 1:3 in 1X PBS before mixing with DH5α 1.0 OD_600_ cell suspensions to attain an ∼1:1 starting cell ratio. 20 µL spots of mixtures were plated on LSLB agar in 24-well plates, allowed to dry, and incubated at 37°C for 5 h. Starting mixtures were also serially diluted and plated on antibiotic-containing selective media to enumerate inhibitor and target strains at the initial time point. Following 5 h, competition spots were resuspended in 1 mL 1X PBS within wells, serially diluted, and plated on antibiotic-containing selective media to enumerate inhibitor and target strains. Colony counts at the initial and 5 h time points allowed for competitive index (C.I.) calculations as follows: C.I. = (inhibitor_*t*5_/target_*t*5_)/(inhibitor_*t*0_/target_*t*0_). A positive log_10_ C.I. indicates the inhibitor strain outcompeted the target strain, a negative log_10_ C.I. indicates the target strain outcompeted the inhibitor strain, and a log_10_ C.I. of ∼0 indicates neither strain had a competitive advantage.

Liquid competitions between *B. cenocepacia*-*P. aeruginosa* co-infection isolates were set up following the above protocol, except 20 µL of cell mixtures were inoculated into 1 mL LSLB and grown for 5 h at 37°C shaking at 220 rpm. For competitions between *Bc*AU1054 and *P. aeruginosa* clinical isolates, *Bc*AU1054 WT and Δ*hcp* strains constitutively expressing *lacZ* were used and inocula/competitions were plated onto antibiotic-containing LSLB agar with 40 µg/mL 5-bromo-4-chloro-3-indolyl β-D-galactopyranoside (X-Gal) to help differentiate between *P. aeruginosa* and *Bc*AU1054 colonies. Competitions between *Bc*AU1054 and pJN-*rsmZ*/pJN105-harboring *P. aeruginosa* teenage/adult isolates were conducted on LSLB agar containing 0.1% L-arabinose.

### *Bc*AU1054 T6SS E/I screen

Co-cultures were set up following the same protocol as in the interbacterial competition experiments. For monocultures, cell suspensions (at an OD_600_ of 1.0) were mixed 1:1 with 1X PBS before plating. For co-cultures and mono-cultures, 20 µL spots were plated on LSLB agar within 24-well plates, spots were allowed to dry, and plates were incubated at 37°C for ∼20 h. Following incubation, the cultures were resuspended in 1 mL 1X PBS within wells, 100 µL were added to 96-well plates, and OD_600_ values and GFP fluorescence intensities (485 nm excitation, 530 nm emission) were measured on a PerkinElmer Wallac VICTOR^3^_™_ plate reader.

### Hcp1 immunoblotting

*P. aeruginosa* strains were swabbed onto LSLB agar and grown overnight at 37°C. For pJN-*rsmZ*/pJN105-harboring *P. aeruginosa* teenage/adult isolates, strains were swabbed onto LSLB agar containing 75 µg/mL gentamicin and 0.1% L-arabinose and grown overnight at 37°C. Following overnight incubation, cells were scraped off plate, resuspended in 1 mL cold 1X PBS, centrifuged for 2 min at 15,000 rpm, washed in 1 mL cold 1X PBS, and diluted to an OD_600_ of 5.0. Cells were then centrifuged for 2 min at 15,000 rpm and resuspended in 200 µL 2X SDS-PAGE sample loading buffer (6X SDS-PAGE sample loading buffer: 375 mM Tris-HCl, 9% sodium dodecyl sulfate (SDS), 50% glycerol, 0.03% bromophenol blue, 1.3 M β-mercaptoethanol), boiled at 99°C for 15 min, and samples were sheared 10 times through a 26G needle. Samples were resolved on 12% SDS-PAGE gels (5 µL loaded), transferred to nitrocellulose membranes, and membranes were blocked with 5% (w/v) non-fat dry milk in 1X PBS for 1 h with rotation at room temperature (RT). Membranes were then washed three times in 1X PBS and incubated with α-Hcp1 peptide antibody (diluted 1:1000 in 5% (w/v) non-fat dry milk in 1X PBS+0.1% Tween®20 (PBS-T)) for 1 h with rotation at RT. Membranes were then washed three times in 1X PBS-T, incubated with IRDye® 800CW-conjugated α-rabbit IgG antibody (diluted 1:25,000 in 5% (w/v) non-fat dry milk in 1X PBS-T) for 30 min with rotation at RT, washed three times in 1X PBS, and imaged on a LI-COR Odyssey® fluorescence imager.

### Sequencing

Genomic DNA was purified from *P. aeruginosa* isolates C078C, C120C, and C123D using the Promega Wizard Genomic DNA Purification Kit. Paired-end TruSeq (Illumina) libraries were generated and sequenced on the Illumina MiSeq 2×150 platform at the High-Throughput Sequencing Facility at the University of North Carolina at Chapel Hill. Demultiplexed FASTQ files were mapped to the PAO1 reference genome (assembly GCA_000006765.1) using the Geneious Prime standard assembler. Sequencing reads can be accessed in BioProject PRJNA609958.

To sequence *P. aeruginosa* CEC isolate genomes, genomic DNA was isolated using a GenElute Bacterial Genomic DNA Kit (Sigma Aldrich, NA2110; St. Louis, MO) following kit instructions with the following exception: all DNA was eluted in 400uL of ultra-pure DEPC-treated water (ThermoFisher Scientific, Waltham, MA). Concentration of DNA preps was determined using a NanoDrop 1000 (ThermoFisher Scientific, Waltham, MA). All preps were stored at −20C. The 150bp sequencing reads from the Illumina platform were assembled using spades v.3.7.1 with careful mismatch correction and the assemblies were filtered to contain only contigs ≥500bp with ≥5X k-mer coverage. The assemblies were further examined for characteristics that would suggest the genome was of high quality (<400 contigs) and potentially *P. aeruginosa*. All reads and assemblies are deposited at NCBI under BioProject PRJNA607994.

Specific *P. aeruginosa* co-infection isolate genes were sequenced by PCR-amplifying genes of interest and submitting the PCR products for Sanger sequencing.

### Bioinformatic analysis of *Bc*AU1054 T6SS-encoding genes and effector proteins

The *Bc*AU1054 and *Bc*J2315 (genome assembly GCA_000009485.1) T6SS-encoding core clusters were aligned in Geneious Prime using the Mauve plugin (Darling *et al*., 2004). Phyre2 (Kelley *et al*., 2015) and HHpred (Zimmermann *et al*., 2018) were used to predict the secondary structures and catalytic activities of potential *Bc*AU1054 T6SS effector proteins.

## ACKNOWLEDGEMENTS

We thank John Mekalanos for graciously providing the α-Hcp1 peptide antibody, and John LiPuma for graciously providing the *B. cenocepacia*-*P. aeruginosa* co-infection isolates. We thank members of the Cotter Lab for helpful discussion while designing experiments and analyzing data for this study. This work was supported by NIH awards NIHR01GM121110 to P.A.C. and U19AI110820 to D.A.R, a graduate stipend from the University of Maryland-Baltimore Graduate School to C.E.C., and Cystic Fibrosis Foundation award COTTER18I0 to P.A.C. Funders had no role in the planning, execution, or analysis of experiments, nor in manuscript preparation and submission.

## SUPPLEMENTAL FIGURES AND TABLES

**Figure S1.**
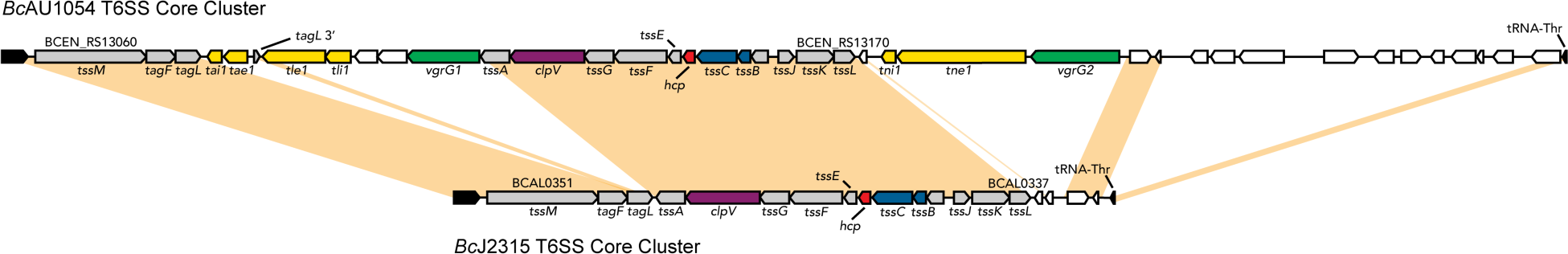
The *Bc*AU1054 T6SS core cluster is homologous to the *Bc*J2315 T6SS core cluster, though contains insertions. The genomic regions encoding the T6SSs of *Bc*AU1054 and *Bc*J2315 were aligned to each other using Mauve. Orange shading represents regions of homology (≥90% nucleotide identity). All the structural protein-encoding genes are homologous (≥90% nucleotide identity) between the two strains, though the *Bc*AU1054 core cluster has multiple insertions – regions spanning *tae1*-*tai1, vgrG1*-*tle1, vgrG2*-*tni2*, and genes downstream of the core cluster predicted to encode proteins involved in phage and mobile genetic elements (see **Figure 1**).

**Figure S2.**
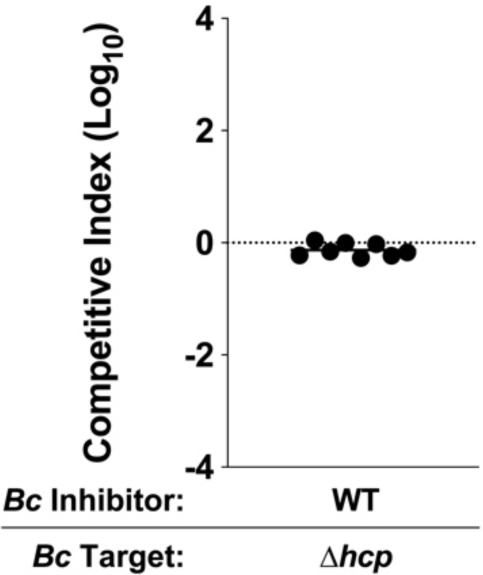
*Bc*AU1054 Δ*hcp* does not have a growth defect. 5 h co-cultures between *Bc*AU1054 WT and Δ*hcp* strains. Circles represent individual co-cultures from two biological replicates, each with four technical replicates. Solid horizontal line represents mean log_10_ C.I. Dotted horizontal line (log_10_ C.I. = 0) indicates no competitive advantage for either inhibitor or target.

**Figure S3.**
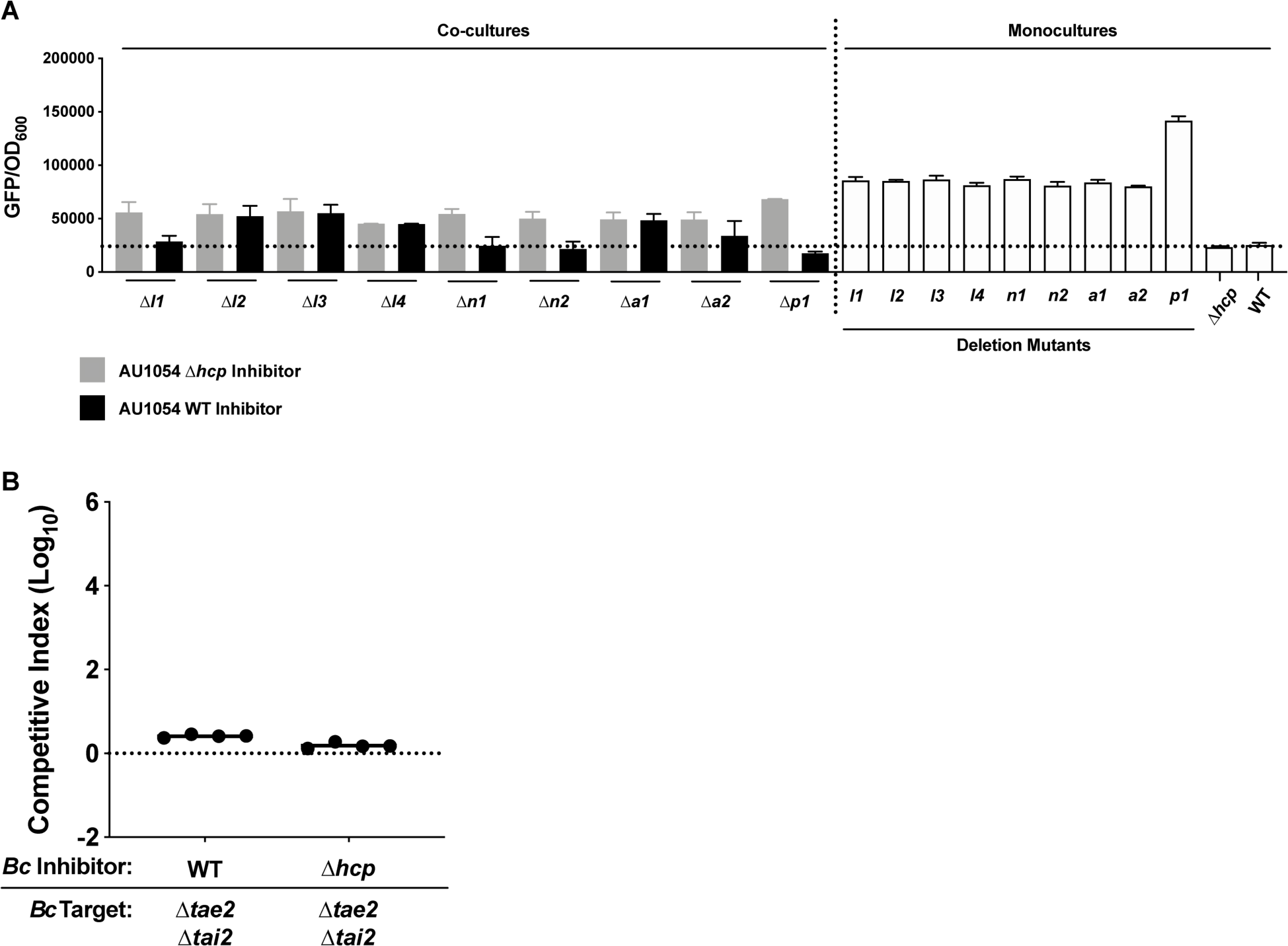
Screening approach to identify probable *Bc*AU1054 T6SS E-I pairs. (A) GFP/OD_600_ values for ∼20 h co-cultures between WT and Δ*hcp Bc*AU1054 inhibitor strains and *Bc*AU1054 E-I deletion mutants constitutively producing GFP, as well as GFP/OD_600_ values for ∼20 h monocultures of each strain. GFP measured at 485 nm excitation and 530 nm emission. Data resulting from at least two biological replicates, with height of bar representing mean GFP/OD_600_ value and error bars representing one standard deviation. Dotted horizontal line represents baseline GFP/OD_600_ value, taken from non-GFP-producing WT and Δ*hcp Bc*AU1054 monocultures. See **Figure 1** and **Table S2** for location of E-I-encoding gene pairs and predicted effector activity, respectively. (B) 5 h interbacterial competition experiment between WT and Δ*hcp Bc*AU1054 inhibitor strains and *Bc*AU1054 Δ*tae2*Δ*tai2* target strain. Circles represent individual co-cultures (four technical replicates for each competition). Solid horizontal lines represent mean log_10_ C.I. values. Dotted horizontal line (log_10_ C.I. = 0) indicates no competitive advantage for either inhibitor or target.

**Figure S4.**
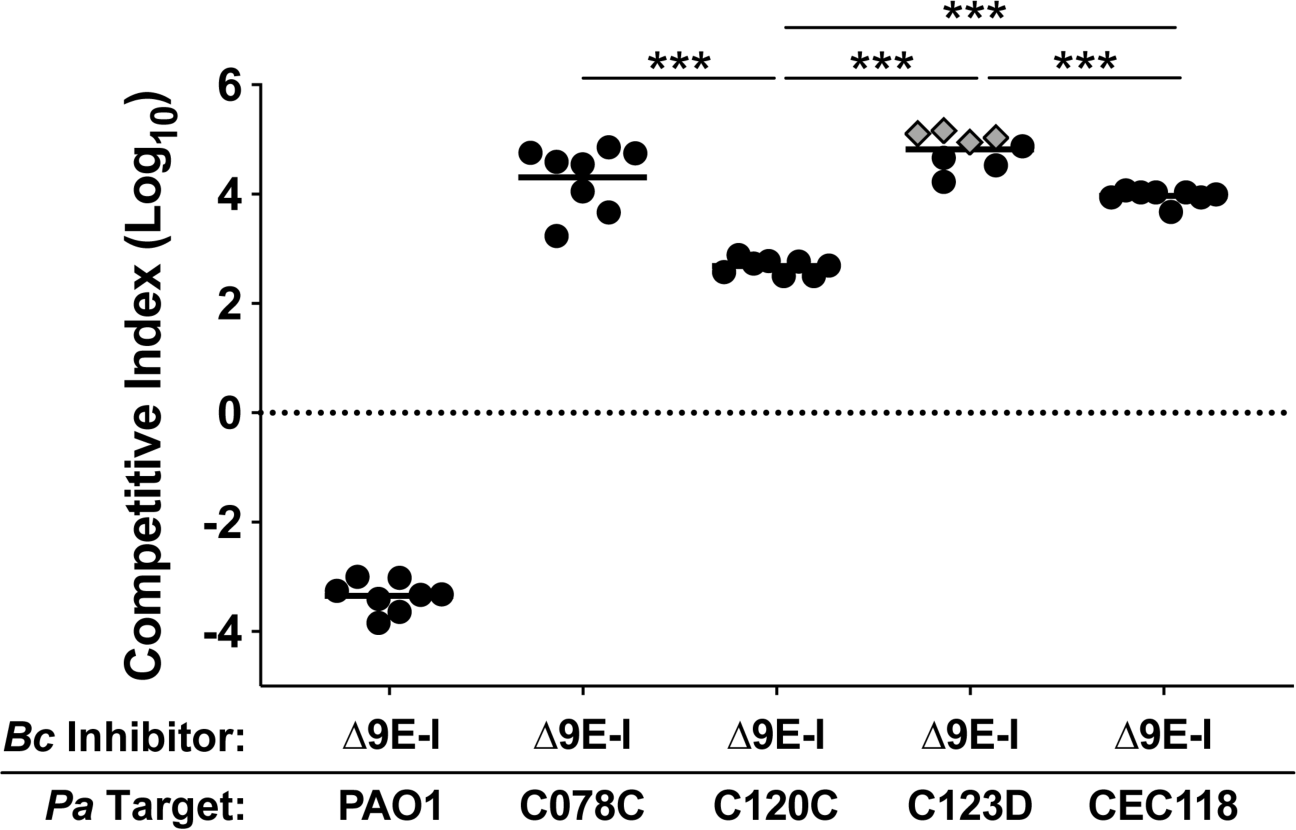
*Bc*AU1054 T6SS effectors exhibit target strain-specific variability in toxicity. Interbacterial competition experiments between *Bc*AU1054 Δ9E-I inhibitor strain and PAO1, C078C, C120C, C123D, and CEC118 *P. aeruginosa* target strains. Circles/diamonds represent individual co-cultures from two biological replicates, each with four technical replicates. Grey-filled diamonds represent competitions from which no target cells were recovered. Solid horizontal lines represent mean log_10_ C.I. values. Dotted horizontal line (log_10_ C.I. = 0) indicates no competitive advantage for either inhibitor or target. \*\*\**P*<0.0005, Mann-Whitney test.

**Table S1. Whole genome sequencing errors in *Bc*AU1054 T6SS genes.**

Correct sequences provided (determined by PCR amplification and Sanger sequencing) for two genes misannotated as pseudogenes – *clpV* and *tne1*. Sequences also provided for *tni1* and *tni2*, which are not annotated.

**Table S2.**
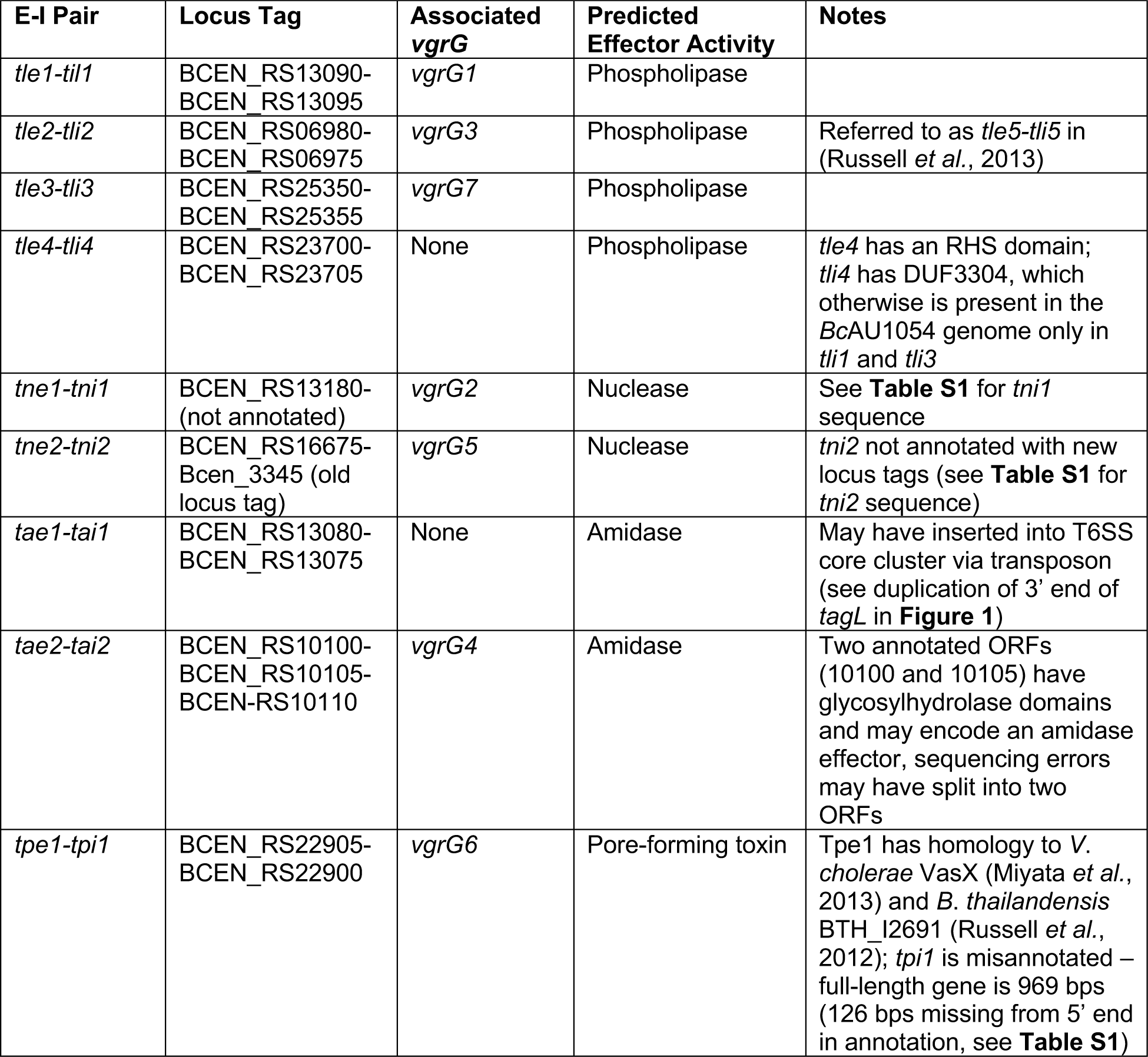
Bioinformatically-predicted *Bc*AU1054 T6SS E-I pairs and predicted effector enzymatic activities.

## Strains Used in this Study

| Strain Name | Source | WGS Identifier |
| --- | --- | --- |
| *Burkholderia cepacia* complex strains |  |  |
| BcAU1054 | BcRLR* | GCA_000014085.1 |
| BcAU1054 $\Delta hcp$ | This paper | |
| BcAU1054 $\Delta hcp attTn7::hcp$ | This paper | |
| BcAU1054 $\Delta tle1\Delta tli1$ | This paper | |
| BcAU1054 $\Delta tle1\Delta tli1 attTn7::tli1$ | This paper | |
| BcAU1054 $\Delta tle2\Delta tli2$ | This paper | |
| BcAU1054 $\Delta tle3\Delta tli3$ | This paper | |
| BcAU1054 $\Delta tle4\Delta tli4$ | This paper | |
| BcAU1054 $\Delta tne1\Delta tni1$ | This paper | |
| BcAU1054 $\Delta tne1\Delta tni1 attTn7::tni1$ | This paper | |
| BcAU1054 $\Delta tne2\Delta tni2$ | This paper | |
| BcAU1054 $\Delta tne2\Delta tni2 attTn7::tni2$ | This paper | |
| BcAU1054 $\Delta tae1\Delta tai1$ | This paper | |
| BcAU1054 $\Delta tae2\Delta tai2$ | This paper | |
| BcAU1054 $\Delta tpe1\Delta tpi1$ | This paper | |
| BcAU1054 $\Delta tpe1\Delta tpi1 attTn7::tpi1$ | This paper | |
| BcAU1054 $\Delta 9E-I$ | This paper | |
| BmCGD2M | BcRLR* | GCA_000182295.1 |
| BmCGD2M *tssC1::pAP82* | This paper |  |
| BdAU0158 | BcRLR* | GCA_000959505.1 |
| BdAU0158 *tssC1::pAP83* | This paper |  |
| BcAU4392 | BcRLR* |  |
| BcAU5161 | BcRLR* |  |
| BcAU7623 | BcRLR* |  |
| BcAU10618 | BcRLR* |  |
| BcAU19695 | BcRLR* |  |
| BcAU22760 | BcRLR* |  |
| BcAU23782 | BcRLR* |  |
| BcAU29704 | BcRLR* |  |
| *Pseudomonas aeruginosa* strains |  |  |
| PAO1 | (Holloway, 1955) Wolfgang Lab collection | GCA_000006765.1 |
| PAO1 *vipA1::Tn* | (Held *et al.*, 2012) |  |
| CEC32 |  | BioProject PRJNA607994 |
| CEC36 |  | BioProject PRJNA607994 |
| CEC42 |  | BioProject PRJNA607994 |
| CEC44 |  | BioProject PRJNA607994 |
| CEC66 |  | BioProject PRJNA607994 |
| CEC73 |  | BioProject PRJNA607994 |
| CEC83 |  | BioProject PRJNA607994 |
| CEC87 |  | BioProject PRJNA607994 |
| CEC112 |  | BioProject PRJNA607994 |
| C078C | (Radlinski *et al.*, 2017) (referred to as BC236) | BioProject PRJNA609958 |
| C078C *attTn7::pppA* | This paper |  |
| C120C | (Radlinski *et al.*, 2017) (referred to as BC238) | BioProject PRJNA609958 |
| C123D | (Radlinski *et al.*, 2017) (referred to as BC239) | BioProject PRJNA609958 |
| CEC118 |  | BioProject PRJNA607994 |
| CEC118 *attTn7::fha1* | This paper |  |
| CEC119 |  | BioProject PRJNA607994 |
| CEC120 |  | BioProject PRJNA607994 |
| CEC121 |  | BioProject PRJNA607994 |
| CEC122 |  | BioProject PRJNA607994 |
| CEC116 |  | BioProject PRJNA607994 |
| *PaAU4391* | BcRLR* |  |
| *PaAU5159* | BcRLR* |  |
| *PaAU7618* | BcRLR* |  |
| *PaAU10617* | BcRLR* |  |
| *PaAU19694* | BcRLR* |  |
| *PaAU22775* | BcRLR* |  |
| *PaAU23781* | BcRLR* |  |
| *PaAU29744* | BcRLR* |  |
| *Escherichia coli* strains |  |  |
| DH5α | Cotter Lab collection | |

\**Burkholderia cepacia* Research Laboratory and Repository, University of Michigan, Ann Arbor, MI USA.

